# Characterization of emerging Oropouche virus tropism and pathogenicity

**DOI:** 10.64898/2026.03.25.714204

**Authors:** Hugo Bruant, Patricia Jeannin, Virginie Geolier, Vincent Mouly, Emeline Perthame, Nassim Mahtal, Jeanne Pascard, François Piumi, Dominique Rousset, Pierre-Emmanuel Ceccaldi, Muriel Coulpier, Valérie Choumet

## Abstract

**Background:** Oropouche virus is an emerging arbovirus increasingly associated with neurological complications, but its human cellular tropism and potential routes to the central nervous system remain poorly defined. This study aimed to characterize infection across clinically relevant human cell types and to investigate interactions with a human blood–brain barrier model and human neuronal/glial cells.

**Methods:** A panel of human cell lines and primary human cells relevant to systemic and neurological disease was infected with Oropouche virus. Viral replication and production of infectious particles were quantified using molecular assays and infectivity titrations, and viral protein expression was assessed by immunoblotting and immunofluorescence. Barrier crossing was evaluated using a Transwell brain endothelial model with permeability monitoring, and infection dynamics in neuronal/glial cultures derived from human neural progenitors were quantified by imaging-based analyses. Group comparisons used non-parametric tests with Dunn–Bonferroni correction and Mann–Whitney tests; neuronal/glial cell counts were analysed using linear models with Fisher tests for interaction terms and multiplicity-adjusted post hoc comparisons.

**Results:** Oropouche virus productively infected hepatocyte-like and intestinal epithelial cells, with high viral RNA output and release of infectious progeny. Primary synoviocytes, chondrocytes and skeletal muscle cells were permissive but produced lower infectious titers. Brain endothelial cells were inoculated and virus was progressively detected in the basolateral compartment, while endothelial permeability remained unchanged, indicating barrier crossing without disruption. In neuronal/glial cultures, both neurons and astrocytes were susceptible; infection was associated with marked cytopathic changes and a preferential, accelerated decline in neuron abundance over time.

**Conclusions:** These findings demonstrate broad human cell tropism and support blood–brain barrier crossing without major loss of barrier integrity, alongside pronounced neuronal vulnerability. The described models provide a platform to dissect mechanisms of neuroinvasion and to evaluate targeted antiviral strategies.

## Introduction

Arthropod-borne viruses (arboviruses) constitute a large and diverse group of viruses transmitted by hematophagous arthropods, including mosquitoes, midges, ticks, and sandflies. These viruses belong to at least 11 distinct viral families, most of which are RNA viruses whose high mutation rates facilitate rapid adaptation to new hosts and ecological niches (1). More than 600 arboviruses have been catalogued by the CDC, and their global relevance has increased due to climate change, urban expansion, and ecosystem disturbances, which collectively increase human exposure to competent vectors (2, 3). In response to accelerating arbovirus emergence, the World Health Organization launched the Global Arbovirus Initiative in 2022 to strengthen surveillance, preparedness, and outbreak response capacities worldwide (4). Orthobunyaviruses represent a medically and economically important genus within the *Peribunyaviridae* family (order Bunyavirales). These enveloped viruses possess a tripartite, negative-sense RNA genome encoding the glycoproteins Gn and Gc, the nucleoprotein N, and the RNA-dependent RNA polymerase L. They also express the nonstructural proteins NSs and NSm, which contribute to innate immune evasion, virulence, and virion assembly (5, 6). Their segmented genome enables frequent reassortment events, generating novel viral variants with altered host range, transmissibility, or pathogenicity (7, 8). Several orthobunyaviruses have caused significant disease outbreaks in humans and animals: Bunyamwera virus (BUNV) induces febrile illness in Africa (9); La Crosse virus (LACV) is a major cause of pediatric encephalitis in the United States (10); Cache Valley virus infects livestock and occasionally humans (11); and Schmallenberg virus (SBV), discovered in 2011, caused widespread epidemics in ruminants across Europe with substantial economic impact (12).

Among orthobunyaviruses, Oropouche virus (OROV) stands out as one of the most clinically important. First isolated in 1955 in Trinidad from a febrile patient (13), OROV has since become a major public health concern in South America, where it has caused more than thirty documented urban epidemics, particularly in the Brazilian Amazon, resulting in over 500,000 infections (14, 15). Its primary vectors include the biting midge *Culicoides paraensis* and the mosquito *Culex quinquefasciatus* (16, 17), although additional vector species likely contribute to transmission cycles. Since 2021, more than 11,000 cases have been reported across South America and the Caribbean, including the first documented fatalities during the 2024 outbreak in Brazil. The recent detection of OROV in French Guiana (18), combined with imported cases in the United States and Europe (19), highlights its potential for geographic expansion beyond endemic regions.

Clinically, OROV infection—Oropouche fever—presents as a dengue-like febrile illness characterized by fever, headache, myalgia, arthralgia, and rash (15). Although typically self-limiting, up to 5–10% of patients develop neurological complications such as meningitis and encephalitis, confirming the virus’s neuroinvasive capacity (20). Other severe manifestations, including hemorrhagic symptoms, persistent neurological sequelae, and fatal outcomes, have been reported in recent outbreaks (19). Vertical transmission has also been documented, leading to miscarriages and congenital anomalies (21), raising concerns about maternal–fetal transmission similar to what is observed with Zika virus (ZIKV) (22). Despite this growing evidence of severe disease, OROV remains significantly understudied compared to other arboviruses in the Americas such as dengue virus, Chikungunya virus (CHIKV), and ZIKV. Major knowledge gaps persist regarding OROV’s cellular and tissue tropism, mechanisms of dissemination within the host, and its ability to interact with and across physiological barriers such as the blood–brain barrier (BBB). Only limited data exist concerning its infection of human neuronal or musculoskeletal tissues, its potential for persistence in immune-privileged sites, or its molecular determinants of neuroinvasion. Addressing these questions is crucial not only for understanding OROV pathogenesis but also for anticipating its epidemic potential, particularly in densely populated areas or regions newly colonized by the vector or the virus.

The objective of this study was therefore to comprehensively investigate the human cellular tropism and neuroinvasive properties of OROV. To achieve this, we evaluated viral replication in a broad panel of human cell types including hepatocytes, epithelial cells, endothelial brain cells, synoviocytes, chondrocytes and skeletal muscle cells to determine tissue susceptibility and identify potential reservoirs of infection. We also employed an established *in vitro* BBB model to assess OROV’s ability to infect and cross a brain endothelial cells monolayer. Finally, we used cultures of human neuronal/glial cells (hNGCs) derived from human neural progenitor cells (hNPCs) to examine direct infection of neurons and astrocytes and quantify viral induced cytopathic effects.

Together, these approaches provide a detailed and multi-dimensional analysis of OROV’s pathogenic potential, clarify its mechanisms of neuroinvasion and cellular injury, and offer critical experimental foundations for future antiviral, diagnostic, and surveillance strategies.

## Materials and Methods

### Cell Culture

Vero E6 cells (ATCC CRL-1586) were used for viral production and titration.

Human enterocyte-like Caco-2 cells (Clone TC7, SCC209, Sigma-Aldrich) and immortalized human hepatocytes HuH-7 (23) were used in infection experiments. All these cells were grown in DMEM medium supplemented with L-glutamine (Gibco, Thermo Fisher Scientific), 10% fetal bovine serum (FBS), 100 U/mL penicillin, and 100 μg/mL streptomycin (P/S), and were maintained at 37 °C and 5% CO_2_.

Human muscle cells (CHQ) were originally isolated from the quadriceps muscle, as previously described (24) and cultivated in Ham’s F-10 supplemented with 50mg/mL of gentamycin and 20% FBS.

Human chondrocytes (HC) and synoviocytes (HFLS) were obtained from Cell Applications, (Ref PB-402-05a and 408K-05a, respectively) and cultured according to manufacturer’s instructions.

Immortalized human brain endothelial cells (hCMEC/D3, Merck, SCC066) were cultivated using endothelial growth medium (EBM-2) (Lonza) supplemented with 5% FBS, 1.5 μM hydrocortisone (Merck), 5 μg/ml ascorbic acid (Merck), 1% chemically defined lipid concentrate (Thermo Fisher Scientific), 1 ng/ml human bFGF (Merck), 10 μM HEPES (Thermo Fisher Scientific), and antibiotics in flask pre-coated with rat collagen-I. Asian tiger mosquito larva cells C6/36 (25) were cultivated in L-15 media (Gibco) supplemented with 10% FBS and 1% P/S.

Human NPCs were originally isolated from the cortex of a first-trimester human fetus, cryopreserved and maintained in continuous cultures as previously described in (26, 27). Differentiation into a mixed population of hNGCs was achieved as previously described (28, 29). Briefly, hNPCs were seeded on matrigel-coated plates at a density of 30,000 cells/cm^2^. Twenty four hours after plating, differentiation was induced by withdrawing EGF (Eurobio) and bFGF (Eurobio) and replacing N2A medium with 1:1 N2A and NBC media (N2A: advanced Dulbecco’s modified Eagle medium-F12 supplemented with 2 mM L-glutamine, 0.1 mg/ml apotransferrin, 25 μg/ml insulin, and 6.3 ng/ml progesterone; NBC: neurobasal medium supplemented with 2 mM L-glutamine and B27 without vitamin A 1X—Invitrogen, Life Technologies). Differentiation conditions were maintained for 13 days with medium replacement twice a week, prior to infection.

### Ethics

Muscle cells were provided by the AFM Tissue Bank (Paris, France). Written informed consent was obtained prior to tissue collection and transfer to MyoBank, a member of the EuroBioBank network. All procedures were conducted in accordance with French bioethics legislation and were approved by the French Ministry for Higher Education and Research (Paris, France) under authorization numbers AC-2013-1869 and AC-2019-3502. For hNPCs, human fetal tissue was collected following legal abortion, with full written informed consent. All procedures for the procurement and use of central nervous system (CNS) tissue were approved and supervised by the Ethics Committee for the Protection of Persons in Biomedical Research at Henri Mondor Hospital, France. HNPCs are officially registered with the French Ministry of Research under the following reference numbers: AC-2017-2993 (“Centre de Ressources Biologiques”, Angers University Hospital, BB-0033-00038) and DC-2019-3771 (UMR Virology, Maisons-Alfort, France).

### Viruses

Oropouche strain Saul/17225/2020 was isolated by Dr D. Rousset in French Guiana (18). The virus was first amplified on C6/36 cells for 3 days at 27°C without CO_2_ and then passed on Vero cells for 3 days at Multiplicity Of Infection (MOI) 0.1 to obtain a working stock. Viral titers were assessed by standard plaque assay on Vero cells using a 0,5% carboxymethyl cellulose (CMC) overlay diluted with DMEM supplemented with 2% FBS. Lysis plaques were then observed visually using Crystal Violet solution.

### Cell infections

Cells were cultivated and infected in 24well plates (Falcon) for infection kinetics and glass coverslips were placed in some wells for immuno-microscopy. hNPCs were differentiated and infected in black 96-well µClear plates (Greiner) for immuno-microscopy. For all infections, virus inoculum was prepared at different MOIs (10, 1, 0.1, 0.01, 0) using the cell type specific culture medium supplemented in 2% FBS. Inoculums were discarded after 1h of incubation at 37°C and replaced with fresh medium supplemented with 2% FBS. Supernatants were then collected at 2h, 24h, 48h and 72h post-infection for analysis.

### Viral RNA production measurements

Parts of cell supernatants were collected and used for RNA extraction using Nucleospin RNA Virus extraction kit (Macherey Nagel). Oropouche S segment was then amplified by RT-qPCR using FNF and FNR primers described by the team of Naveca *et al*. (30). RT-qPCR reactions were prepared using the Power SYBR^tm^ Green RNA-to C_T_ ^tm^ 1-Step Kit (Applied Biosystems) in 20µl volume and performed using CFX96 Touch Real-Time PCR Detection System (Bio-Rad). Qiagen digital PCR (dPCR) system was also used for absolute quantification using the same primers. The dPCR were realized using QIAcuity OneStep Advanced EG Kit (Qiagen).

### Titration using xCELLigence technology

Agilent xCELLigence technology was used for viral titration of infected cells supernatant. Titration were realized on Vero cells using RTCA E-Plates (Agilent), as previously described (31). Cells were seeded 24h before infection and incubated at 37°C, 5% CO_2_. Basal impedance is then measured using the RTCA analyzer xCELLigence RTCA DP (Agilent) for 6 days. This allows to measure the time needed for 50% degradation of the cell monolayer (T_D50_). Before samples analysis, virus dilutions were titrated by standard plaque assay before being measured by RTCA to obtain a standard curve correlating titer and T_D50_. Supernatant were then put on cells and Cell Index were normalized at t=0.

### Western Blot

At different time post-infection, cells were lysed using RIPA Buffer (ThermoFischer) coupled with Protease Inhibitor (Roche). Western Blot analysis was then performed on the cell lysates for detection of viral protein. Lysates were mixed with Nupage LDS Sample buffer (Invitrogen) and put at 70°C for protein denaturation. Sample proteins were separated by migration in a NuPage 4-12% Bis-Tris Gel (Invitrogen) and transferred on a 0,2 µm thick Polyvinylidene fluoride (PVDF) membrane (Biorad). Oropouche virus nucleoprotein (OROV N) was detected using a mouse polyclonal antibody kindly provided by Dr. B. Brennan’s team from Glasgow University. An anti-calnexin (CNX) polyclonal rabbit antibody (ADI-SPA-865; Enzo Life Sciences) was also used for control. Revelation was performed using secondary anti-mouse HRP-conjugated antibodies and Pierce ECL Western Blotting Substrate (Thermo Fisher Scientific).

### Immunomicroscopy

Infected cells were fixed using 4% paraformaldehyde during 45 min at room temperature. Cells were then permeabilized and saturated at the same time using a bovine serum albumin (BSA) 3%, Triton 0.3%, PBS 1X solution for 2h at room temperature under agitation. Primary antibodies were then diluted in PBS with BSA 0.3%, Triton 0.03% and incubated overnight at 4°C. Oropouche virus immunoreactivity was assessed using a murine monoclonal anti-Gc antibody (1:50 diluted) (EVAg, 022A-05675) kindly provided by Dr. S. Reiche’s Team from Friedrich-Loeffler-Institut. For hNPCs, rabbit polyclonal or mouse monoclonal anti-ß-III-Tubulin antibodies (1:1000 diluted) (Sigma, T2200, T8660 respectively) were used for neuron immunoreactivity assays. Astrocyte immunoreactivity was determined using monoclonal rabbit anti-GFAP (Glial Fibrillary Acidic Protein) antibodies (1:1000 diluted) (Sigma, G3893). Cells were washed 3 times using PBS solution. Secondary antibodies such as Alexa Fluor 488-conjugated AffiniPure Goat Anti-Mouse IgG (H+L) (Jackson ImmunoResearch, 115-545-003) and Cyanin 5-conjugated AffiniPure Donkey Anti-Rabbit IgG (H+L) (Jackson ImmunoResearch, 711-175-152) were diluted at 1:500 in the same solution as the primary antibodies mentioned above and incubated for two hours at room temperature under agitation in the dark. Cells were then washed 3 times with PBS. For hNPCs, cells were then incubated with PBS, 0.1 µg/mL DAPI (Sigma, D9542) during 45 min at room temperature under agitation in the dark. Cells were finally washed once with PBS before observation. For other cell types, washed glass coverslips were taken out of wells and placed on microscope slides with ProLong® Gold Antifade with DAPI (Molecular Probes, Cell Signaling Technology).

### Image analysis

For quantification of hNPCs in 96-well plates, images were acquired with an Opera Phenix Plus (Revvity) running Harmony v5.23, in epifluorescence mode, with a 10x/NA 0.3 (quantification) and 40x/NA 1.1 objective (imaging) with a binning of two. The fluorescent channels were selected as follows: DAPI (excitation/emission filter wavelengths (Exc/Em), in nm): 405/435-480; Alexa 488: 488/500-550; Alexa 568: 565/570-630; Alexa 647: 633/650-760. Nine fields were acquired per well and merged for image quantification and only 4 fields for imaging. All images were transferred and analysed using Signals Image Artist software (Revvity, v1.4.2). Nuclei were segmented and only complete nuclei, not touching the edges of the image, were processed. Alexa 488 intensities, reflecting infection, were determined within a slightly larger nuclear region (two pixels larger than each nucleus). Potential dead cells were excluded by thresholding on the DAPI channel, as apoptotic/dead cells have brighter nuclei. The remaining cells were classified according to their size to differentiate neurons from astrocytes, the latter being considered to have a nucleus > 100 µm². Finally, a positivity threshold was determined by evaluating the single-cell infection-related signal distributions in the positive and negative control populations to discriminate infected cells, and it was applied to all conditions and cell types. On average, ∼32,000 cells were quantified per well.

For image acquisition, hNGCs were imaged of the different other cell types infected, cells were imaged using an EVOS microscope (AMG) using AMEP 4698, Plan Fluor 20x/0.5 objective. The fluorescent channels were selected as follows: DAPI: 405/435-480 and Alexa 488: 488/500-550.

### Statistical models for cell count analysis

Cell counts obtained from images of labeled hNGCs were used to develop linear models. For each dose, a linear model was adjusted to the logarithm of the number of cells. The model included the effects of time, as well as its interaction with cell type and experimental repetition. The significance of the interaction term was assessed using a Fisher’s test, with p-values adjusted using Bonferroni correction. The percentage of variance explained by each effect was determined from the ANOVA table, using the variance decomposition property of the model. Post hoc comparisons between doses and durations were performed using the R package ‘emmeans: Estimated Marginal Means, aka Least-Squares Means v1.11.1’, with p-values adjusted using the ‘mvt’ correction procedure.

### Blood-Brain Barrier experiments

The *in vitro* model of BBB used was previously described (32, 33). Human endothelial cells hCMEC/D3 were grown on Transwell Clear-TM filters (polyester, 12mm diameter, pore size 3μm; BD Biosciences) in EBM-2 (Lonza) without hydrocortisone. At confluence, hydrocortisone was added, and cells formed tight junctions when cultured for at least 6 days. BBB integrity was assessed by FITC-Dextran (70 kDa; Merck) permeability as previously described (32). In brief, hCMEC/D3 endothelial monolayers grown on Transwell inserts were transferred into culture wells containing 1.5 mL of RPMI (Gibco, Thermo Fisher Scientific) medium without phenol red. The medium within the inserts was substituted with 500 μL of RPMI containing an equivalent concentration of FITC Dextran. The cells were incubated at 37°C for 10 minutes. Subsequently, the inserts were sequentially relocated to new wells filled with 1.5 mL of fresh RPMI and incubated at 37°C for 15 and 20 minutes, respectively. Quantification of FITC Dextran concentration in the basolateral compartments was carried out using a fluorescence spectrophotometer (TriStar2 LB 942, Berthold), with fluorescence emission detected at 535 nm following excitation at 485 nm. The apparent endothelial permeability coefficient (Pe) of FITC Dextran was calculated in cm/min as outlined in a previously established protocol (34).

## Results

### Susceptibility of Vero E6, HuH-7 and Caco-2 cell lines to OROV

Vero E6, HuH-7, Caco-2 cell lines were inoculated with OROV at different MOIs (0.1; 1 and 10) and viral replication and production were assessed using various methods. Detection of viral RNA by dPCR at 24, 48, 72 hours post infection (p.i.) revealed a significant increase in RNA levels over time, indicating viral RNA production in the three cell lines (Fig. 1A shows the results obtained at a MOI 0.1). In addition, we assessed viral nucleoprotein production within the cells by Western blotting at different times post-infection (Fig. 1B). The figure shows presence of OROV N protein in infected cells at 24h p.i. for Vero and HuH7 cells and at 48h p.i. for Caco-2 cells, suggesting an effective translation of viral RNA. These cell lines were able to produce infectious viruses, at 24, 48 and 72h p.i., ranging between 10^4^ and 10^8^ PFU/mL (Fig. 1C). Eventually, the different cell lines were also immunoreactive for the viral glycoprotein Gc (see Fig. 1D) which seems to be concentrated in the perinuclear region.

**Figure 1:**
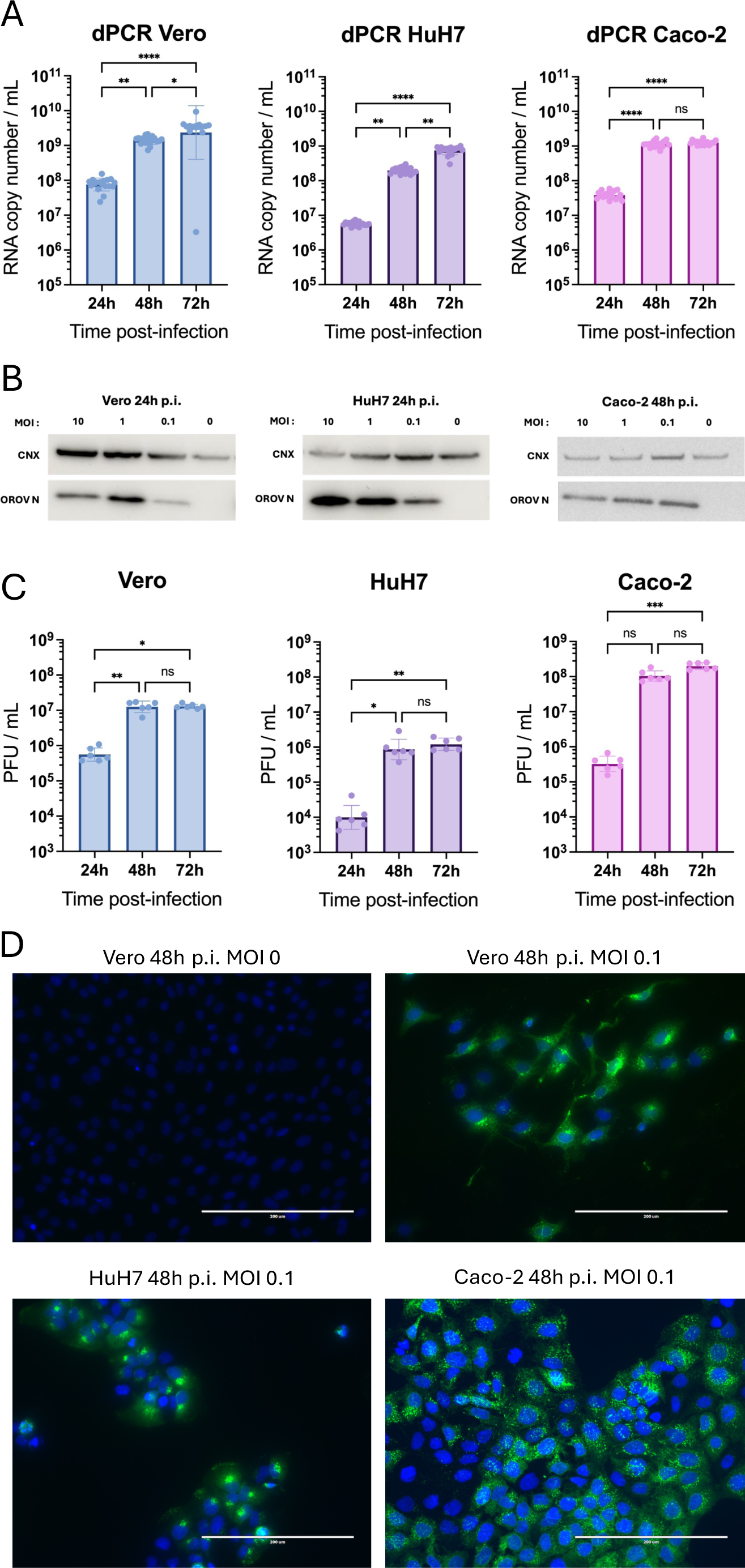
Susceptibility of different cell lines to OROV. A: dPCR results of Vero (left), HuH7 (middle) and Caco-2 (right) cells supernatants infected with OROV at MOI 0.1 and collected at different times (24, 48, 72 h p.i.). The graphs show 18 points per condition as well as the mean and geometric standard deviation. ns: not significant; *: p-value < 0.05; **: p-value < 0.01; ***: p-value < 0.001; ****: p-value < 0.0001 (Dunn-Bonferroni non-parametric test). B: Vero (left), HuH7 (middle), and Caco-2 (right) cells were infected with OROV at different MOIs. Cell lysates were collected at 24 or 48 h p.i. and analysed by Western blot, using antibodies against calnexin (CNX) as internal control and against the N protein of OROV (OROV N). C: The supernatants from Vero (left), HuH7 (middle) and Caco-2 (right) cultures were collected at different times after infection with OROV at MOI 0.1. The viral titer was determined on Vero cells using xCELLigence technology from a standard range. Each condition has 6 replicates. ns: not significant; *: p-value < 0.05; **: p-value < 0.01; ***: p-value < 0.001; ****: p-value < 0.0001 (Dunn-Bonferroni non-parametric test). D: Vero (upper right picture), HuH7 (left, down) and Caco-2 (right, down) cells were infected with OROV at MOI 0.1. Upper left: mock-infected. At 48 h p.i., OROV Gc protein was detected (green) and nuclei were counterstained (blue). The scale bar represents 200 µm.

### Susceptibility of human primary cells to OROV

Based on the clinical manifestations associated with OROV infections, different primary human cell cultures were selected for their physiological relevance *in vitro*. Human myoblasts (CHQ), chondrocytes (HC) and synoviocytes (HFLS) were used to model muscle and joint damage. Cultures were inoculated with OROV at different MOIs. Viral RNA production could be detected in chondrocytes, synoviocytes and myoblasts at MOI 0.1, at 24, 48 and 72 h p.i. (Fig. 2A). Western blot analysis after labelling the N protein in cell lysates indicated a reduced level of N protein expression in the different cells (chondrocytes, synoviocytes, myoblasts), as shown in Fig. 2B. Low viral production, increasing between 24 and 72 h. p.i., was observed for synoviocytes, while a reduced number of infectious particles is detected for chondrocytes, meanwhile infection of myoblasts shows no evidence of increased viral production over time (Fig. 2C).

**Figure 2:**
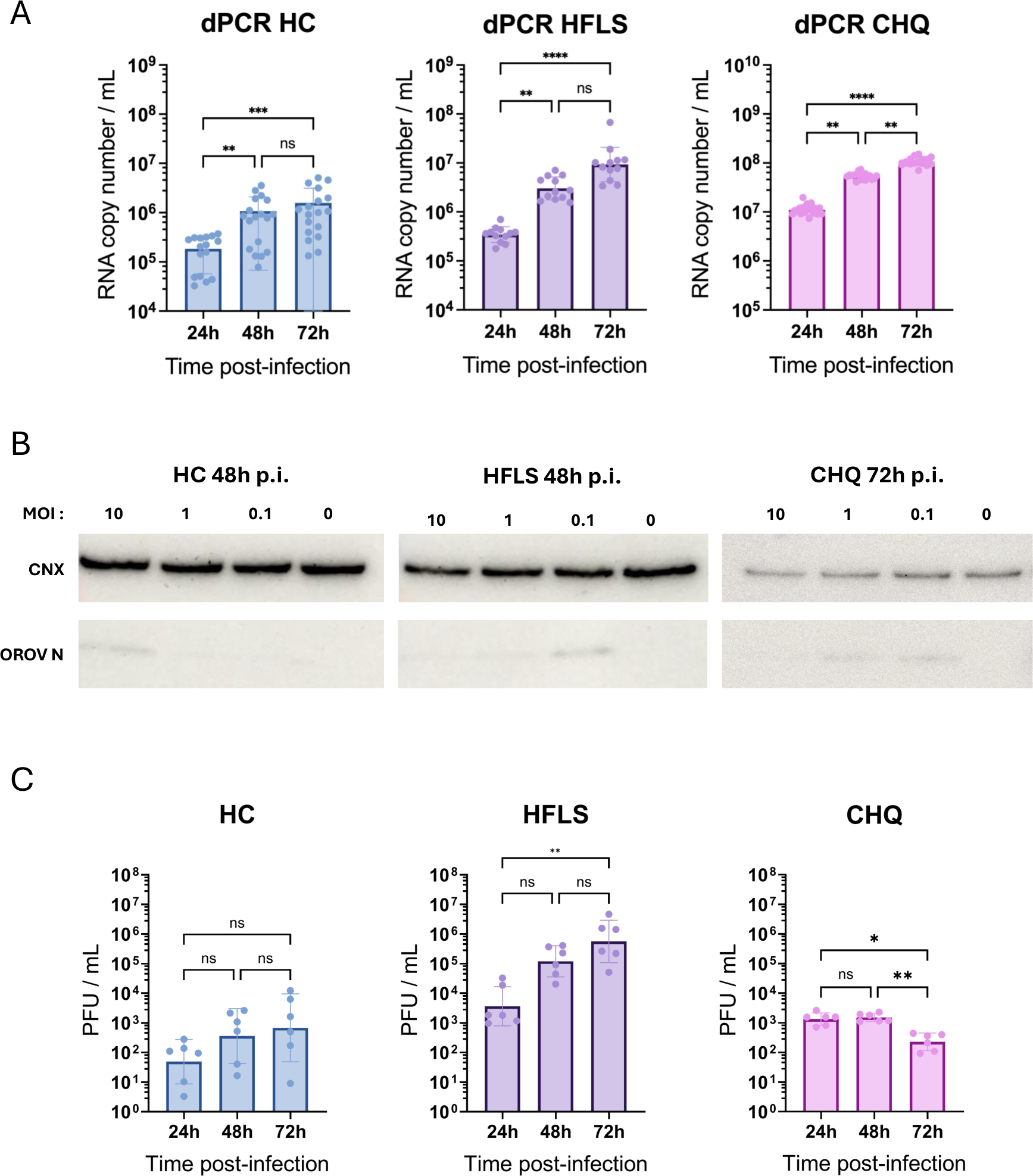
Susceptibility of different human primary cells to OROV. A: dPCR results of human chondrocytes (HC, left), Fibroblast-Like Synoviocytes (HFLS, middle) and muscular cells (CHQ; right) cultures supernatants infected with OROV at MOI 0.1 and collected at different times (24, 48, 72 h p.i.). The graphs show 18 points (except HFLS with 12 points) per condition, as well as the mean and geometric standard deviation. ns: not significant; *: p-value < 0.05; **: p-value < 0.01; ***: p-value < 0.001; ****: p-value < 0.0001 (Dunn-Bonferroni non-parametric test). B: HC (left), HFLS (middle), and CHQ (right) cells were infected with OROV at different MOIs. Cell lysates were collected at 48 or 72h p.i. and analysed by Western blot, using antibodies against calnexin (CNX) as internal control and against the N protein of OROV (OROV N). C: The supernatants from HC (left), HFLS (middle) and CHQ (right) cultures were collected at different times after infection with OROV at MOI 0.1. The viral titer was determined on Vero cells using xCELLigence technology from a standard range. Each condition has 6 replicates. ns: not significant; *: p-value < 0.05; **: p-value < 0.01 (Dunn-Bonferroni non-parametric test).

In immunofluorescence, few cells were immunoreactive for the viral Gc protein at 24 h p.i. for HFLS and CHQ, but at 48 h p.i. immunoreactive cells were observed for all cell types, as well as 72 h. p.i. (Fig. 3)

**Figure 3:**
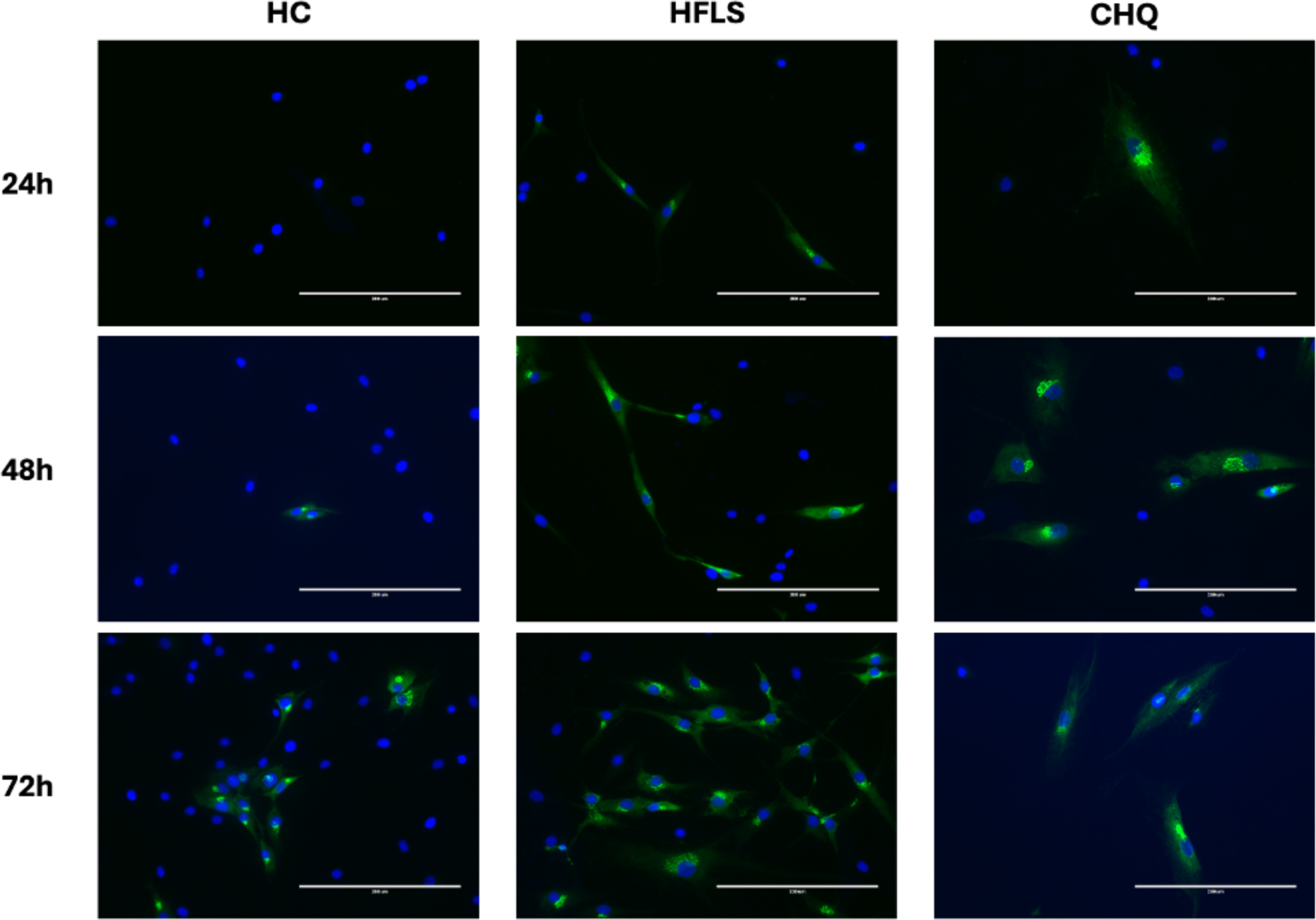
Immunolabelling of the OROV Gc protein in different primary human cells. HC, HFLS and CHQ cells were infected on coverslips with OROV at MOI 0.1. Gc protein of the OROV virus was detected (green), and nuclei were also labelled (blue). The scale bar represents 200 µm.

### Crossing of the BBB by OROV

Since OROV has been shown to be neuroinvasive during epidemics, we investigated the crossing of the BBB by OROV, using an *in vitro* model of BBB (35). Human brain endothelial cells (hCMEC/D3) were grown and differentiated on coverslips or Transwell filters according to the experiments, in order to constitute an endothelial monolayer. Infection of the endothelial cells could be observed by immunofluorescence using anti-Gc antibodies, with an increased signal after 72 h p.i. (Fig. 4).

**Figure 4:**
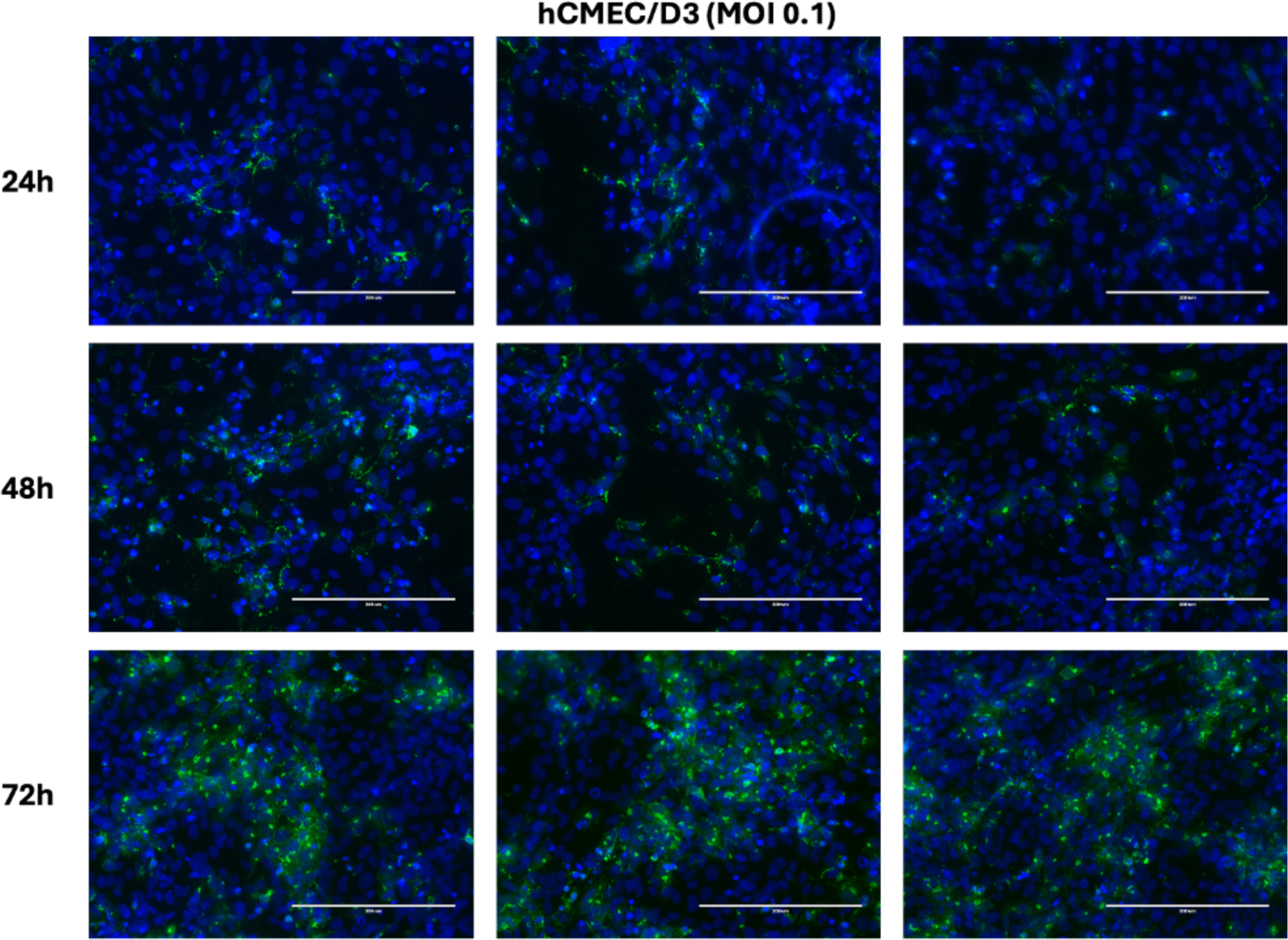
Immunofluorescence analysis of hCMEC/D3 cells. Confluent hCMEC/D3 cells were inoculated with OROV at a MOI of 0.1 on glass coverslips. OROV Gc protein (green) and nuclei (blue) were labelled. The scale bar represents 200 µm.

To assess endothelial barrier infection and evaluate the crossing of the human brain endothelium, cells were also grown and differentiated on Transwell filters, delineating apical and basal compartments. The tightness of the barrier can be measured by permeability assays for fluorescent dyes such as FITC-Dextran (70 kDa). Endothelial cells were inoculated at a MOI 1 in the apical compartment, and viral RNAs were measured both in apical and basal compartments. As shown in Fig. 5A, viral RNAs were detected in both compartments, with a regular increase between 24 and 72 h p.i.. Western blot analyses revealed that the viral nucleoprotein N could be detected in endothelial cells at 48 h p.i. at different MOIs: 0.1, 1 and 10 (Fig. 5B). Yielding of infectious virus particles was demonstrated in both apical and basal compartments, with an increase from 24 h p.i. to 72 h p.i. (Fig. 5C). Interestingly, paracellular passage of viral particles could be excluded since the value of FITC Dextran permeability did not change significantly at the different timepoints between infected and non-infected (Fig. 5D).

**Figure 5:**
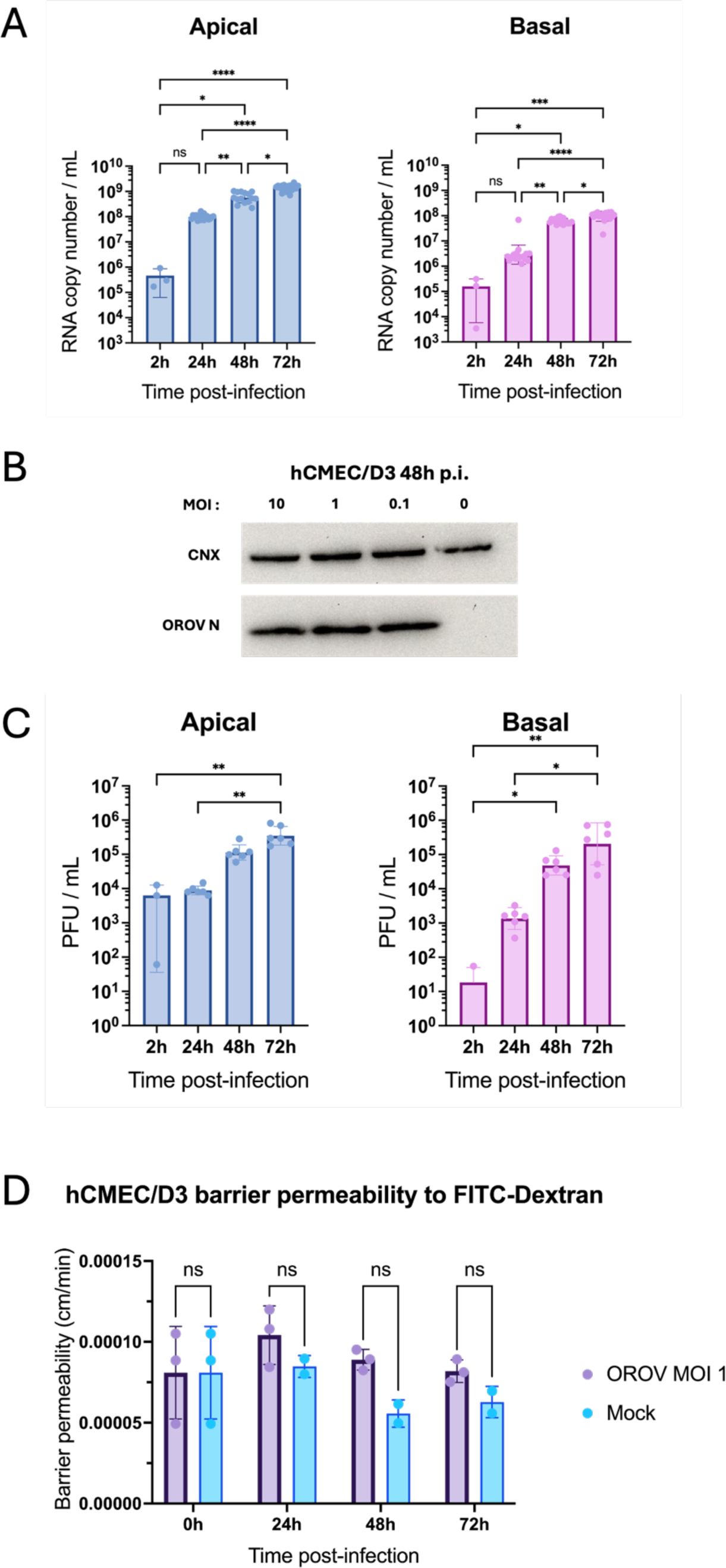
Crossing of an *in vitro* model of BBB by OROV. Experiments were performed on a monolayer of human brain endothelial cells grown on Transwell filters. OROV was inoculated at the apical side (MOI 1). A: The RNAs from the supernatants of the apical and basal compartments were quantified by dPCR at different times p.i.. ns: not significant; *: p-value < 0.05; **: p-value < 0.01; ***: p-value < 0.001; ****: p-value < 0.0001 (Dunn-Bonferroni non-parametric test). B: hCMEC/D3 cells were inoculated with OROV at different MOIs. Cell lysates were collected at 48 p.i. and analysed by Western blot, using antibodies against CNX as internal control and against the N protein of OROV (OROV N). C: The supernatants from hCMEC/D3 cultures were collected in apical and basal compartments at different times after inoculation with OROV at MOI 1. The viral titer was determined on Vero cells using xCELLigence technology from a standard range. Each condition has 6 replicates. ns: not significant; *: p-value < 0.05; **: p-value < 0.01 (Dunn-Bonferroni non-parametric test). D: The integrity of the barrier was assessed by measuring permeability to FITC-Dextran (70kDa). Transwell filters containing monolayers of endothelial cells were placed in new plates and 500µL of FITC-Dextran diluted in culture medium were deposited at the apical level. Fluorescence emission on the basal side was then measured at different timepoints. ns: not significant; *: p-value < 0.05; **: p-value < 0.01; ***: p-value < 0.001; ****: p-value < 0.0001 (2way ANOVA).

### Susceptibility of human neural cells

Given the encephalitis and meningitis symptoms observed during OROV infections (20), we investigated its neurotropism in primary human neuronal/glial cell cultures. At 13 days of differentiation, hNGCs were inoculated with OROV at MOI 0.1 and then analysed over time (24, 72 and 96h p.i.). We first examined whether the virus could productively infect the cells. Viral RNA production was quantified in the supernatant (Fig. 6A), showing an increase from 24 to 72 and 96h p.i.. The presence of infectious viral particles in the supernatant, which peaked at 72h p.i., was further shown by titration assay (Fig. 6B). We next sought to determine what cellular type was susceptible to the virus. Examination of cells stained for either ß-III-Tubulin and viral Gc antibodies (marking neurons) or GFAP and viral Gc antibodies (marking astrocytes) revealed that both cell types were susceptible to OROV infection (Fig. 6C). At 72h p.i., the number of cells appeared considerably reduced. These results and phenotypic modifications such as reduced arborizations were also observed at MOI 1 48 and 72h p.i., compared to non-infected control (MOI 0) (Supplementary Fig. 1).

**Figure 6:**
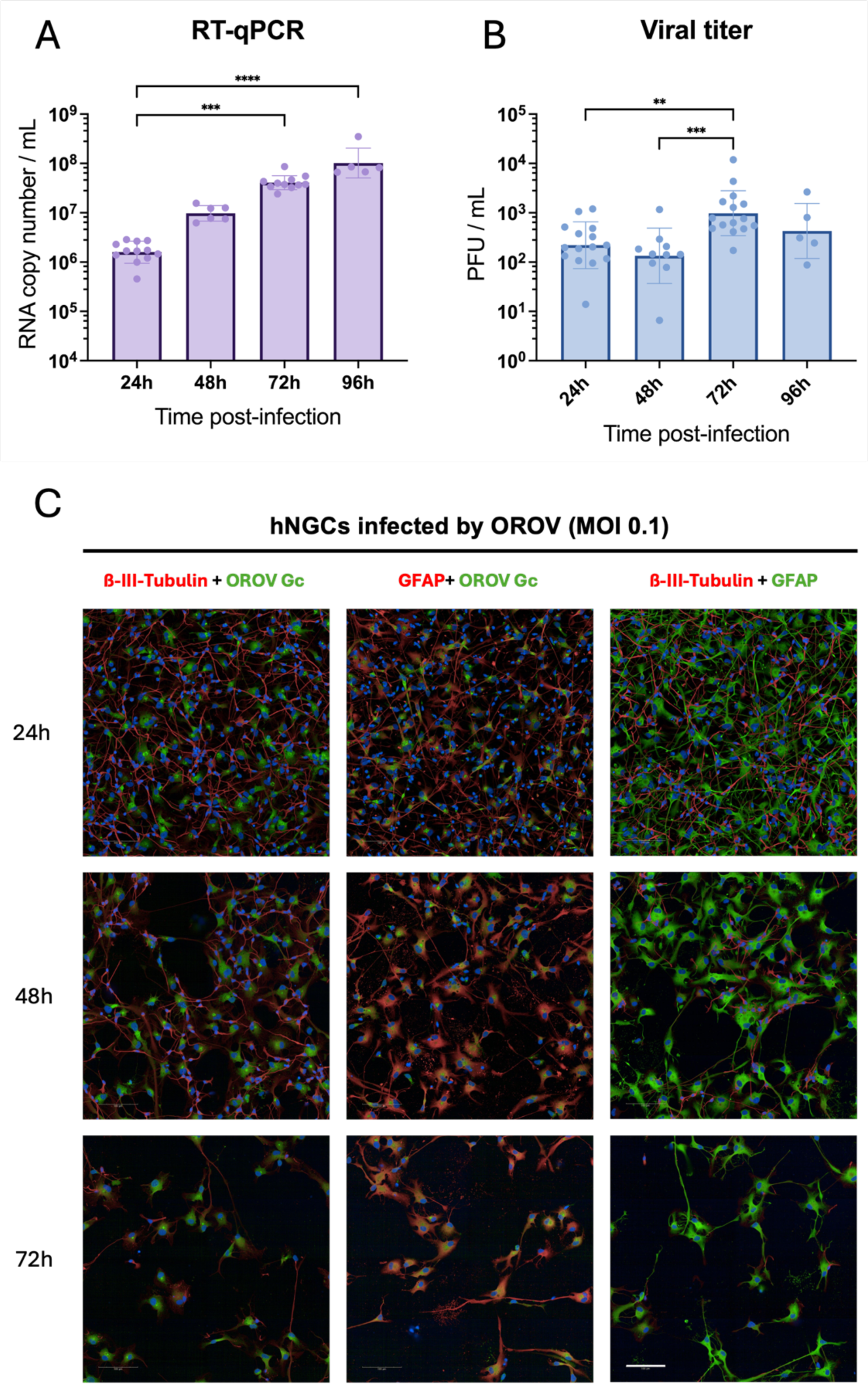
Human neuronal/glial cells susceptibility to OROV infection. Supernatants of hNGCs inoculated with OROV at MOI 0.1 were collected at 24, 48, 72 and 96 h.p.i. and analysed by RT-qPCR (A) or titration assays on Vero cells by xCELLigence (B). *: p-value < 0.05; **: p-value < 0.01; ***: p-value < 0.001; ****: p-value < 0.0001 (Dunn-Bonferroni non-parametric test). C: Immunolabelling with anti-Gc (OROV), anti-ß-III-Tubulin (neurons) and/or anti-GFAP (astrocytes) antibodies, with counterstained nuclei (blue). Scale Bar represents: 100µm.

To confirm that OROV caused marked cellular damage and to assess its specific impact on neurons and astrocytes, we quantified the proportion of these two cell types in differentiated hNGCs cells infected or not with OROV. Quantification of Gc immunoreactivity was then used to distinguish infected from non-infected cells. Statistical analyses based on linear models revealed significant temporal shifts in cell-type proportions (p < 0.001). Absolute cell counts showed a general reduction in total cell numbers in infected cultures (Fig. 7). Both neurons and astrocytes were affected, but neurons declined significantly faster (p < 0.001). Neuronal decay followed an exponential pattern, while astrocytes decreased more linearly (Fig. 7A). Neurons were equal to or more abundant than astrocytes at 24h p.i., depending on MOI (Fig. 7B). Their proportion then decreased to approximatively 25% by 72h p.i. across all MOIs tested. However, no significant difference was observed regarding the susceptibility of both cell types to OROV infection.

**Figure 7:**
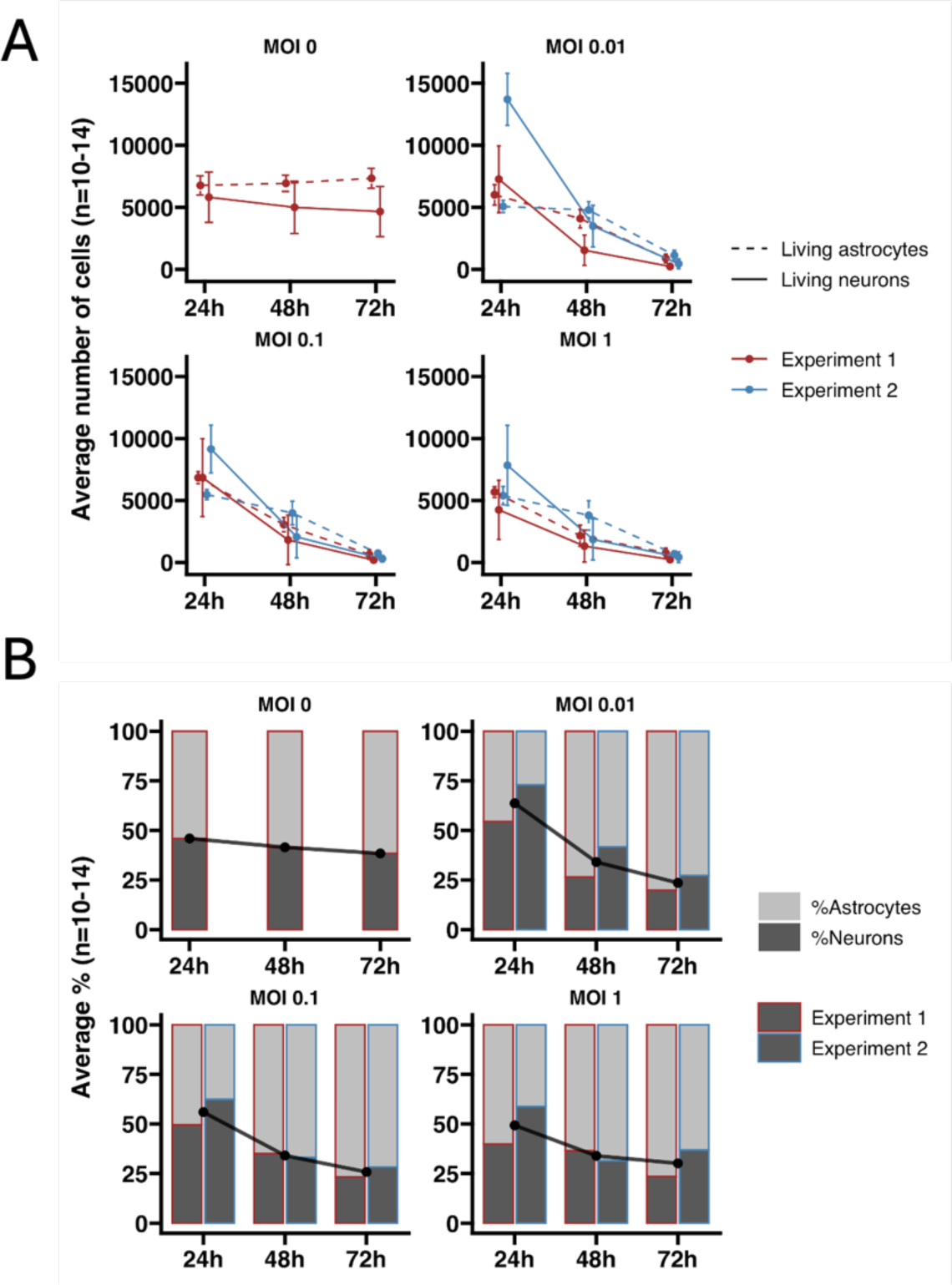
Temporal dynamics of neuronal and astrocytic populations following OROV infection. hNGCs at 13 days of differentiation were infected with OROV at MOI 0, 0.01, 0.1 and 1 for 24, 48 and 72 hours. Cells were distinguished as neurons or astrocytes based on their nuclear size. A: The graphs display the total number of living cells belonging to each cell type. Colors denote the different repetitions of the experiment. Neurons are represented with a solid line, and astrocytes with a dashed line. Dots indicate the average number of cells per experiment and by cell type. Error bars show the 95% confidence interval of the mean. B: The graphs display the proportion of each cell type (neurons and astrocytes) among all viable cells in the wells (n = 10–14 wells). Border colors differentiate between the two repetitions of the experiment, and the fill color of the bar indicates the cell type. The solid black line represents the average across the two experiments.

## Discussion

Oropouche virus (OROV) is an emerging arbovirus of major public health concern. First isolated in Trinidad in 1955, OROV has since caused more than thirty large outbreaks in the Brazilian Amazon, resulting in over half a million cases, and has expanded across South and Central America, including French Guiana (36–38). The 2024 epidemic revealed previously unreported fatal outcomes, as well as maternal–fetal transmission leading to miscarriage and neonatal malformations. Reports of meningitis and encephalitis in infected individuals further suggest that OROV may be neurotropic and neuroinvasive (39–41).

In this study, we aimed to better characterize the cellular tropism and neuroinvasive properties of OROV. By combining a broad *in vitro* analysis of viral replication across diverse human cell types with models of the BBB and neural cell susceptibility assays, we demonstrate that OROV exhibits extensive cell tropism, can cross endothelial barriers, and induces marked cytopathic effects on CNS-derived cells. These data broaden significantly the limited pathophysiological studies available to date.

We used human cell types that reflect clinical features or suspected targets of OROV infection. Hepatocyte-like Huh-7 cells were chosen due to the hepatic dysfunction described in severe human cases (42) and hepatocyte infection documented in hamsters (43). Human epithelial cells (Caco-2) and Vero cells were included to model infection at the vector-bite entry site and potential mucosal transmission, as described for other arboviruses (44, 45). Moreover, study of the epithelial cell line Caco-2 may refer to the intestinal symptoms such as diarrhea that are observed during Oropouche infection. However, immortalized lines such as Caco-2 and Vero cells may have altered receptor expression or growth adaptations that limit their physiological relevance. To complement these models, we used primary human synoviocytes, chondrocytes and skeletal muscle cells, relative to the reported myalgias and arthralgias during OROV infection (39, 46), although, we cannot exclude the implication of other resident cell types such as motor or sensitive neurons. These models are consistent with studies of other arthritogenic arboviruses such as Ross River, Mayaro and CHIKV (47–55), and provide highly relevant human systems for investigating musculoskeletal pathology. Neurotropism was assessed using human neural progenitor cells (hNPCs) differentiated into neurons and astrocytes, a model previously used for Borna disease virus (26), Tick-Borne Encephalitis Virus (TBEV) (28) and West Nile Virus (WNV) (56). This system is well suited to study neuroinvasion, although it does not replicate the complex crosstalk between neural, glial and immune cells as in the *in vivo* situation. Therefore, comparison with murine or hamster infection models (43, 57) and *ex vivo* human brain slices (58) remains essential. Finally, we assessed the neuroinvasive capacity of the virus using the hCMEC/D3 human BBB model (35), a well-established endothelial system validated in more than 650 studies, including work on retroviruses (32, 59, 60) and arboviruses such as dengue virus, ZIKV and Japanese encephalitis virus (JEV) (33, 61–64). Although none of the *in vitro* BBB model fully recapitulates the complexity of the *in vivo* barrier—including immune surveillance and astrocyte–pericyte interactions—this model offers a robust platform to evaluate viral entry, replication and translocation across human brain endothelial cells.

Our findings confirm and expand previous observations that OROV infects a wide range of human cells, including hepatocytes, epithelial cells, endothelial cells, neurons and astrocytes (39, 40). Huh-7 hepatocytes showed robust viral replication, in agreement with recent work confirming their vulnerability and suitability for antiviral screening (65). Importantly, we show for the first time that OROV efficiently infects primary human musculoskeletal cells including synoviocytes, chondrocytes and muscle cells. These results parallel findings from other arboviruses: CHIKV (50, 55), Mayaro (48), Ross River virus (47), Sindbis virus (53), and ZIKV (54) that infect musculoskeletal tissues either *in vitro* and/or *in vivo*. Persistent infection has been demonstrated in chondrocytes for up to 36 days post-inoculation (50), supporting the hypothesis that joint might serve as reservoir and contribute to OROV-associated arthralgia. Although the viral production did not meet the high viral titers of cell lines, this musculoskeletal tropism is consistent with clinical symptoms (39, 46, 66) and suggests that OROV may share pathogenic mechanisms with other arthritogenic viruses. In addition, in neural mixed cultures derived from hNPCs, we provide direct evidence that OROV infects both human neurons and astrocytes. Neurons were highly susceptible, as suggested in another model in a preprint (67), showing robust infection and pron ounced cytopathic effects, supporting a role for direct neuronal injury in the neurological manifestations of OROV infection (40, 68). These findings are consistent with the detection of viral antigen in neurons of infected hamsters (43) and in human brain slices (58), and complement the recent observation of neuronal infection in humans (67). Astrocytes were also highly permissive to infection. However, unlike other neurotropic arboviruses such as ZIKV, WNV, TBEV and JEV which typically induce astrocytic activation characterized by hypertrophy and GFAP upregulation (28, 69–75), OROV infection resulted in reduced GFAP expression and loss of astrocytic arborization, without classical signs of astrogliosis. This distinct behavior suggests a unique pathogenic strategy, whereby neurons may constitute the primary targets of acute injury, whereas astrocytes could sustain long-lasting infection, modulate neuroinflammation, and potentially impact BBB homeostasis. The detection of viral antigen in both cell types, together with the progressive decline in neuronal and astrocyte numbers, echoes observations made for other neurotropic arboviruses such as ZIKV and LACV, where infection of glial and neuronal cells has been associated with encephalitic disease (26, 27, 76). Interestingly, in the same hNPC-derived model, TBEV was also shown to infect both neurons and astrocytes (28), although cytopathic effects appeared more rapidly with OROV in our study. A key difference, however, is the preferential neuronal tropism typically observed for TBEV and other flaviviruses such as JEV (77–79), whereas OROV infects neurons and astrocytes with comparable efficiency. This pattern aligns with some studies reporting preferential infection of neurons and microglia in human brain slices (58) and neurons in experimentally infected suckling mice (57), but together, our results establish OROV as a neurotropic orthobunyavirus with a broader neural tropism and distinct glial responses compared with other encephalitic arboviruses.

Using the hCMEC/D3 model (35), we demonstrate that OROV can infect human brain endothelial cells and cross the BBB without compromising barrier integrity, as indicated by unchanged FITC dextran permeability. Viral RNA and infectious particles were readily detected basolaterally, supporting productive transcellular crossing. These findings align with neuroinvasion pathways described for Rift Valley fever virus (80, 81), ZIKV (63, 64), TBEV (82) and LACV (81), and provide a mechanistic basis for the CNS involvement observed in human OROV cases (39–41). In contrast, the flavivirus JEV was unable to infect human cerebral endothelial cells in an *in vitro* model of BBB (33). Alternative routes, including transcytosis (83–85), Trojan horse mechanisms involving infected leukocytes (86–88), and neural propagation as shown for WNV (89, 90), cannot be excluded and will require further investigation using animal and/or microfluidic BBB models.

Our findings open several avenues for future research. First, the identification of multiple human cell types permissive to OROV provides relevant models to explore viral entry mechanisms and to screen antiviral compounds. The susceptibility of synoviocytes, chondrocytes, muscle cells and astrocytes also raises the possibility of viral persistence in specific tissues, promoting targeted in vivo studies. Establishing mouse models and human brain organoids will be essential to confirm neuroinvasion pathways and CNS pathogenesis. Finally, dissecting the roles of NSs and NSm proteins in our models may reveal key determinants of virulence and new therapeutic targets.

## Acknowledgments

The authors are most grateful to Dr Odile Blanchet (Centre de Ressources Biologiques, BB-0033-00038, CHU Angers, Angers, France) for providing us with the hNPCs, to Cristina Barsan, Renaud Du Pasquier and Juan Pablo Parada Zelada (Qiagen) for helpful advice, Dr Anne Danckaert (UTechS PBI, Pasteur Institut) for the development of the image analysis pipeline and Tatiana Traboulsi (SC Biomarks) for help in dPCR experiments. We also gratefully acknowledge the support of the Photonic BioImaging platform (UTechS PBI) supported by the French National Research Agency (France BioImaging, ANR-24-INBS-0005 FBI (BIOGEN); Investments for the Future), the Institut Pasteur and the Région Île-de-France (DIM1Health program) funding for the use of the Opera Phenix system.

