## Supplemental Data 1 for "Characterization of emerging Oropouche virus tropism and pathogenicity"

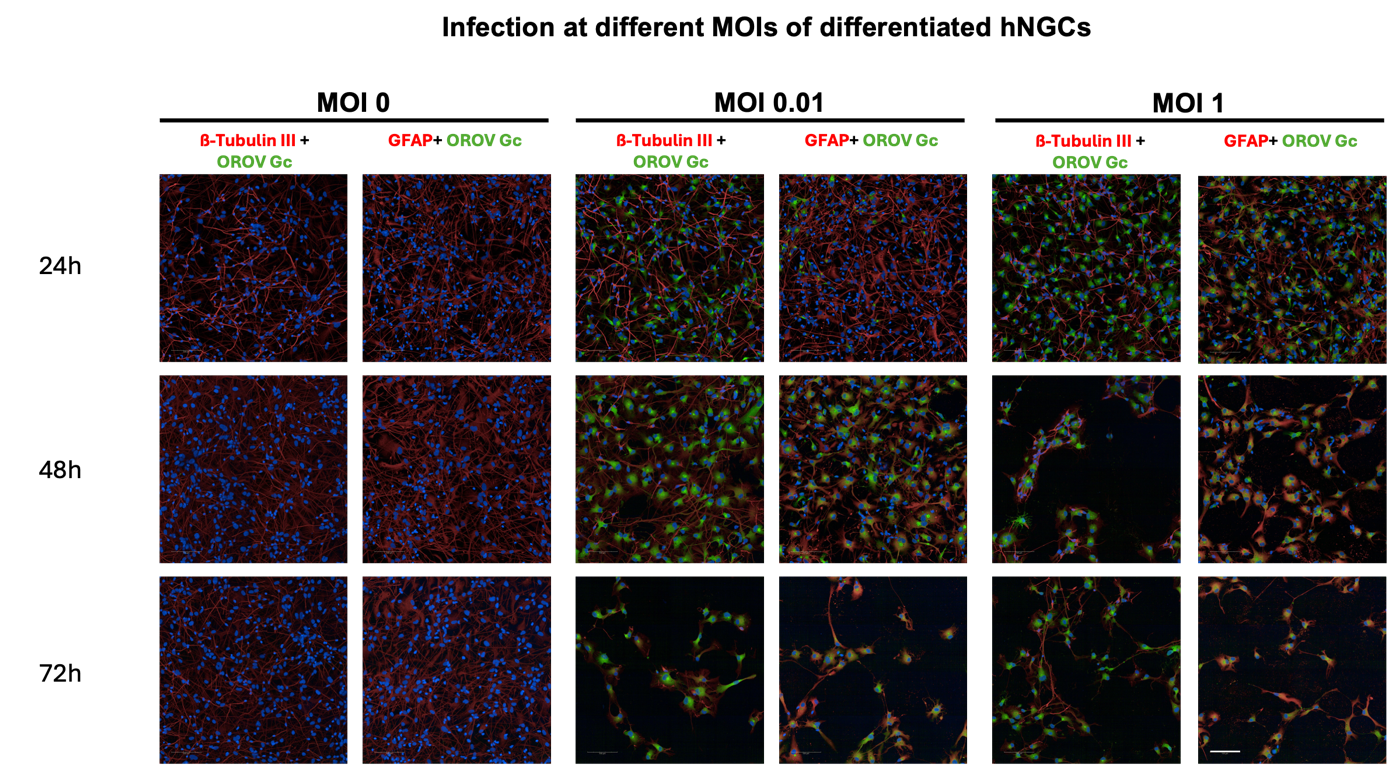
 **Supplementary figure 1 :** Immunolabelling with anti-Gc (OROV), anti-ß-III-Tubulin and/or anti-GFAP antibodies. Scale Bar represents: 100µm.
